# Benchmarking Agentic Bioinformatics Systems for Complex Protein-Set Retrieval: A Coccolithophore Calcification Case Study

**DOI:** 10.64898/2026.03.28.715041

**Authors:** Xiaoyu Zhang

**Affiliations:** California State University San Marcos, San Marcos, California, USA

**Keywords:** agentic AI, bioinformatics benchmarking, UniProt retrieval, coccolithophore, calcification, protein annotation

## Abstract

Large language model agents are increasingly used for bioinformatics tasks that require external databases, tool use, and long multi-step retrieval workflows. However, practical evaluation of these systems remains limited, especially for prompts whose target set is both large and biologically heterogeneous. Here, I benchmarked three agent systems on the same difficult retrieval task: downloading coccolithophore calcification-related proteins from UniProt across six mechanistically distinct categories, while producing category-separated FASTA files and supporting evidence. The compared systems were Codex app agents extended with Claude Scientific Skills, Biomni Lab online, and DeerFlow 2 with default skills only. Outputs were normalized at the UniProt accession level and compared category by category using overlap analysis, Venn decomposition, and a heuristic relevance assessment of each subset relative to the benchmark prompt. Across the six shared categories, Codex retrieved 2,118 proteins, DeerFlow 6,255, and Biomni 8,752 in a run. Codex showed the best balance between sensitivity and specificity: 92.4% of its proteins fell into subsets labeled high relevance and the remaining 7.6% into medium relevance. DeerFlow was substantially more exhaustive, but 43.8% of its proteins fell into low or low-medium relevance subsets. Biomni produced the largest sets, yet 69.5% of its proteins fell into low or low-medium relevance subsets, mainly due to broad expansion into generic calcium sensors, kinases, transcription factors, and poorly specific domain families. Category-specific analysis showed that Codex was the strongest primary source for inorganic carbon transport, calcium and pH regulation, vesicle trafficking, and signaling, whereas DeerFlow contributed valuable complementary matrix and polysaccharide candidates. A second run for each system also separated them strongly by repeatability: Codex had the highest within-system stability (mean category Jaccard 0.982; micro-Jaccard 0.974), DeerFlow was intermediate (0.795; 0.571), and Biomni was least stable (0.412; 0.319). These results suggest that for complex protein-family retrieval tasks, agent quality depends less on raw output volume than on prompt decomposition, taxonomic scoping, exact query generation, provenance-rich export artifacts, and repeated-run stability.

## 1 Introduction

Retrieving proteins from UniProt for a narrowly defined molecular process is straightforward when the target is a single enzyme family or pathway. The problem becomes harder when the biological process is distributed across transport, matrix biology, glycan remodeling, vesicle trafficking, and regulatory signaling, because the desired set is no longer a single ontology term or curated pathway. Coccolithophore calcification is a good example of this difficulty. Coccolithophores produce calcite scales intracellularly inside a Golgi-derived coccolith vesicle, and the process depends on coordnated inorganic carbon acquisition, calcium transport, proton handling, organic matrix assembly, polysaccharide metabolism, vesicle biogenesis, and regulatory control [1–6].

Recent biological studies have sharpened the protein landscape underlying coccolithogenesis. Transcriptomic and gene-expression work in *Emiliania huxleyi* implicated bicarbonate transporters, CAX-like calcium exchangers, vacuolar ATPases, and GPA-associated pathways [3, 7]. A voltage-gated proton channel was shown to support pH homeostasis during calcification [4]. More recent proteomic and matrix studies identified bicarbonate transporters, beta-carbonic anhydrase, coccolith-associated matrix proteins, fibronectin- and MAM-domain proteins, pentapeptide repeats, and carbohydrate-remodeling enzymes as plausible calcification candidates [5, 6]. These signals make coccolithophore calcification biologically rich, but they also make exhaustive retrieval difficult because the relevant annotation space is fragmented and unevenly curated in UniProt.

Agentic AI systems are a plausible solution because they can translate a biologically structured prompt into database queries, code, evidence summaries, and deliverables. The three systems examined here represent different design philosophies. Codex is a general multi-agent desktop environment designed for parallel agent work and skill-based extensions [8]. Claude Scientific Skills is an open skill library that adds domain-specific scientific workflows, databases, and analysis tools to agentic coding systems including Codex [9]. Biomni is a biomedical AI agent centered on tool use, retrieval-augmented planning, and code execution for biomedical research [10, 11]. DeerFlow 2 is an open-source super-agent harness built around skills, sub-agents, memory, and sandboxes [12].

Here, I compare these three systems on a realistic benchmark prompt: download all UniProt proteins related to calcification in coccolithophores, split into separate FASTA files for six required categories, and provide evidence for the selected proteins. The goal was not to measure model latency or token cost, because only output folders were available, but to assess practical retrieval quality. I also used a second repeated run for each system to estimate within-system variance. Specifically, I asked four questions. First, how much do the agents agree on the shared core of calcification-related proteins? Second, how much of each agent’s expansion appears biologically relevant to the prompt rather than generic annotation broadening? Third, how stable are the exported protein sets across repeated prompts within each system? Fourth, what workflow best balances sensitivity, specificity, and repeatability for future AI-assisted bioinformatics retrieval tasks?

## 2 Methods

### 2.1 Benchmark task

The benchmark prompt asked each system to download coccolithophore calcification-related proteins from UniProt and to organize them into separate FASTA files for six biologically motivated categories: (1) inorganic carbon acquisition and carbonate chemistry, (2) calcium delivery and proton or pH homeostasis, (3) organic matrix, crystal templating, and adhesion, (4) matrix-polysaccharide biosynthesis and remodeling, (5) coccolith-vesicle biogenesis, trafficking, and membrane remodeling, and (6) signaling and gene-control regulators. The prompt explicitly named representative protein types in each category, requested exhaustive search within coccolithophores only, and required a separate evidence file.

### 2.2 Agent systems compared

The primary cross-agent comparison used three output folders: `codex_results/run_2,Biomni_lab_output/run_2,` and `deer_flow_output/run_2.` A repeated-run comparison then used the corresponding `run_1` folders for each system. The Codex condition used the OpenAI Codex app with the open-source Claude Scientific Skills library available in the local environment [8, 9]. The Biomni condition used Biomni Lab online, whereas the public Biomni paper and repository were used here only to describe the system at a high level [10, 11]. The DeerFlow condition used DeerFlow 2 with default skills only [12]. Because the benchmark was performed retrospectively on saved outputs, the comparison captures practical behavior on completed runs rather than fully instrumented live sessions. The overall analysis workflow is summarized in Figure 1.

**Figure 1:**
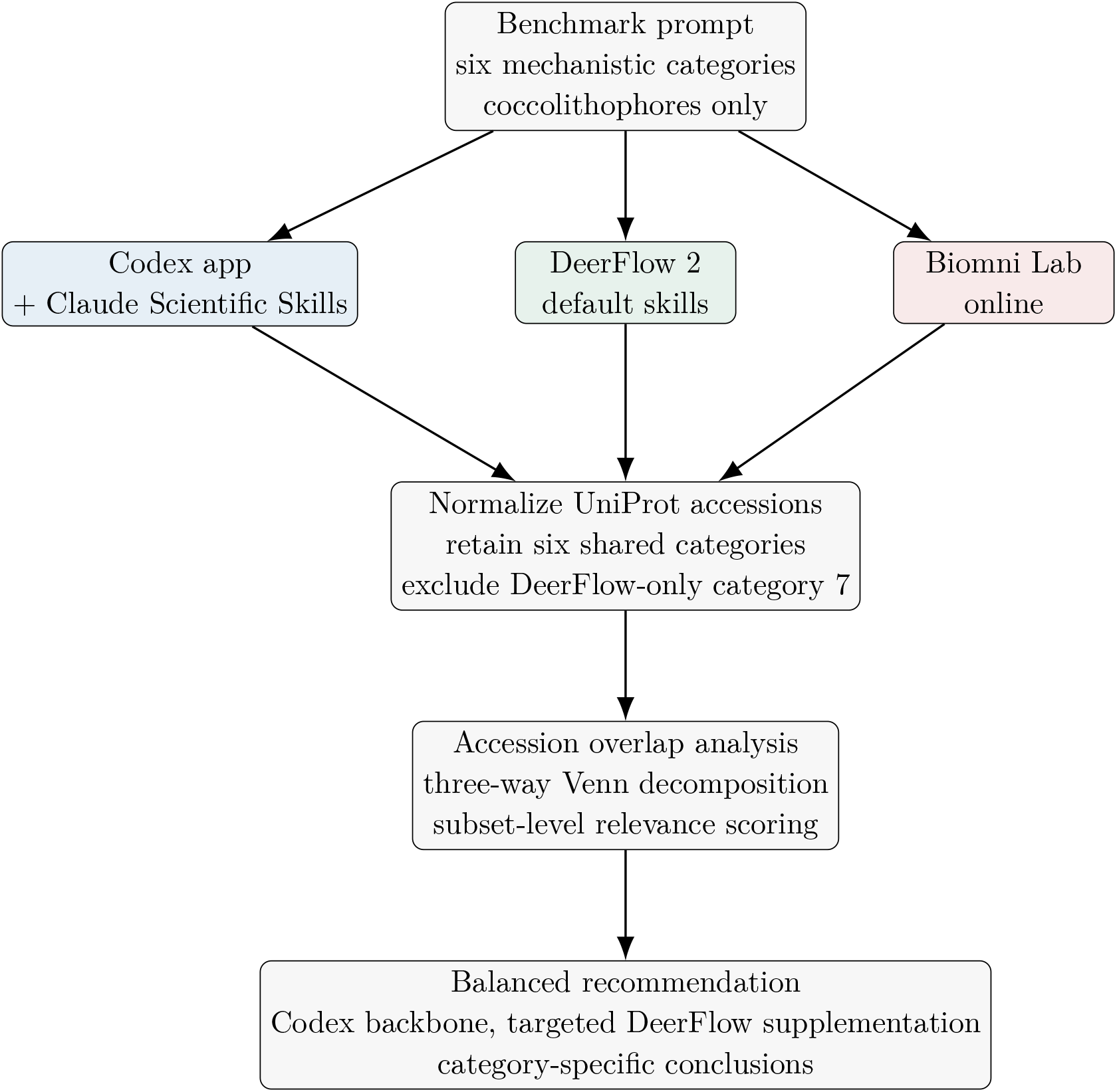
Benchmark workflow used in this study. Saved outputs from the three agent systems were normalized to UniProt accessions, reduced to the six shared prompt categories, and compared using overlap analysis plus a literature-informed relevance heuristic.

### 2.3 Normalization and overlap analysis

All outputs were reduced to the six categories shared by all three systems. DeerFlow produced an additional seventh category related to ion-transport functions; this category was excluded from the primary cross-agent comparison because it was not shared by the other systems and was not required by the prompt. FASTA headers and metadata tables were normalized to UniProt accession sets, and category-specific three-way overlap was computed accession by accession. The resulting report included one Venn decomposition per category and tables for all seven possible set regions (Codex only, DeerFlow only, Biomni only, the three pairwise overlaps, and the triple overlap).

### 2.4 Subset relevance assessment

Each Venn subset was evaluated against the benchmark prompt using a heuristic relevance label. Protein names and descriptions in each subset were summarized into dominant family terms, then classified as *High, Medium, Low-Medium*, or *Low* relevance according to the extent to which they matched the prompt’s expected families and the coccolithophore calcification literature [3–6, 13, 14]. For example, carbonic anhydrases, bicarbonate transporters, sodium/calcium exchangers, V-type proton ATPases, fibronectin- and lectin-domain matrix proteins, glycosyltransferases, sulfotrans-ferases, SNAREs, Rabs, calcineurin-like phosphatases, phospholipase C, and 14-3-3 proteins were treated as positive evidence. Conversely, subsets dominated by aquaporins, generic EF-hand proteins, broad kinase families, generic transcription factors, or many uncharacterized proteins were penalized. This relevance score is a retrieval-quality proxy, not a substitute for manual curation or experimental validation.

### 2.5 Quantitative summary metrics

For each category, I recorded the number of proteins retrieved by each system, the triple-overlap count, and the qualitative source of divergence. For each agent overall, I summed proteins across the six shared categories and calculated the fraction falling into *High*+*Medium* relevance subsets as a specificity proxy. A higher value indicates tighter alignment to the prompt, whereas a larger total protein count indicates broader retrieval sensitivity.

### 2.6 Repeated-run within-system variance

To relax the original single-run limitation, I also compared `run_1` and `run_2` within each agent system across the same six shared categories. FASTA headers were parsed directly to accession sets, and each agent-category pair was compared using intersection size, union size, Jaccard similarity, and directional recovery of `run_1` by `run_2` and vice versa. Because DeerFlow and Biomni changed some category filenames between runs, these files were manually mapped onto the same six canonical category slugs before comparison. The same keyword-based heuristic was then applied to the `run_1`-only, shared, and `run_2`-only subsets in order to distinguish stable cores from run-specific broadening. Aggregate repeatability was summarized both as the mean category Jaccard and as a micro-Jaccard across all category-accession pairs.

### 2.7 Benchmark artifacts

The agents differed not only in FASTA output size but also in provenance artifacts. Codex produced a retrieval script, a test file, a manifest, and accession-level evidence tables. DeerFlow produced per-category TSV files, a combined deduplicated union, a structured `summary.json` with explicit UniProt query terms, and a retrieval script. Biomni produced per-category FASTA files, a large aggregate CSV, a narrative evidence document, and an execution-trace notebook. These differences matter because they affect reproducibility and the ease of downstream manual curation.

## 3 Results

### 3.1 The benchmark captured a clear sensitivity-specificity trade-off

Across the six shared categories, Codex retrieved 2,118 proteins, DeerFlow 6,255, and Biomni 8,752. Figure 2 shows the overall trade-off between total retrieved proteins and the fraction assigned to *High*+*Medium* relevance subsets. Codex occupied the most favorable corner of this comparison, pairing a moderate total with perfect alignment under the heuristic scoring scheme: 1,958 proteins (92.4%) were in high-relevance subsets and 160 (7.6%) in medium-relevance subsets. DeerFlow expanded coverage substantially but lost specificity, with 3,514 proteins (56.2%) in high-relevance subsets, 1,868 (29.9%) in low-medium subsets, and 873 (14.0%) in low subsets. Biomni produced the largest collections but the weakest specificity profile, with only 30.5% of proteins in high- or medium-relevance subsets and 69.5% in low or low-medium subsets.

**Figure 2:**
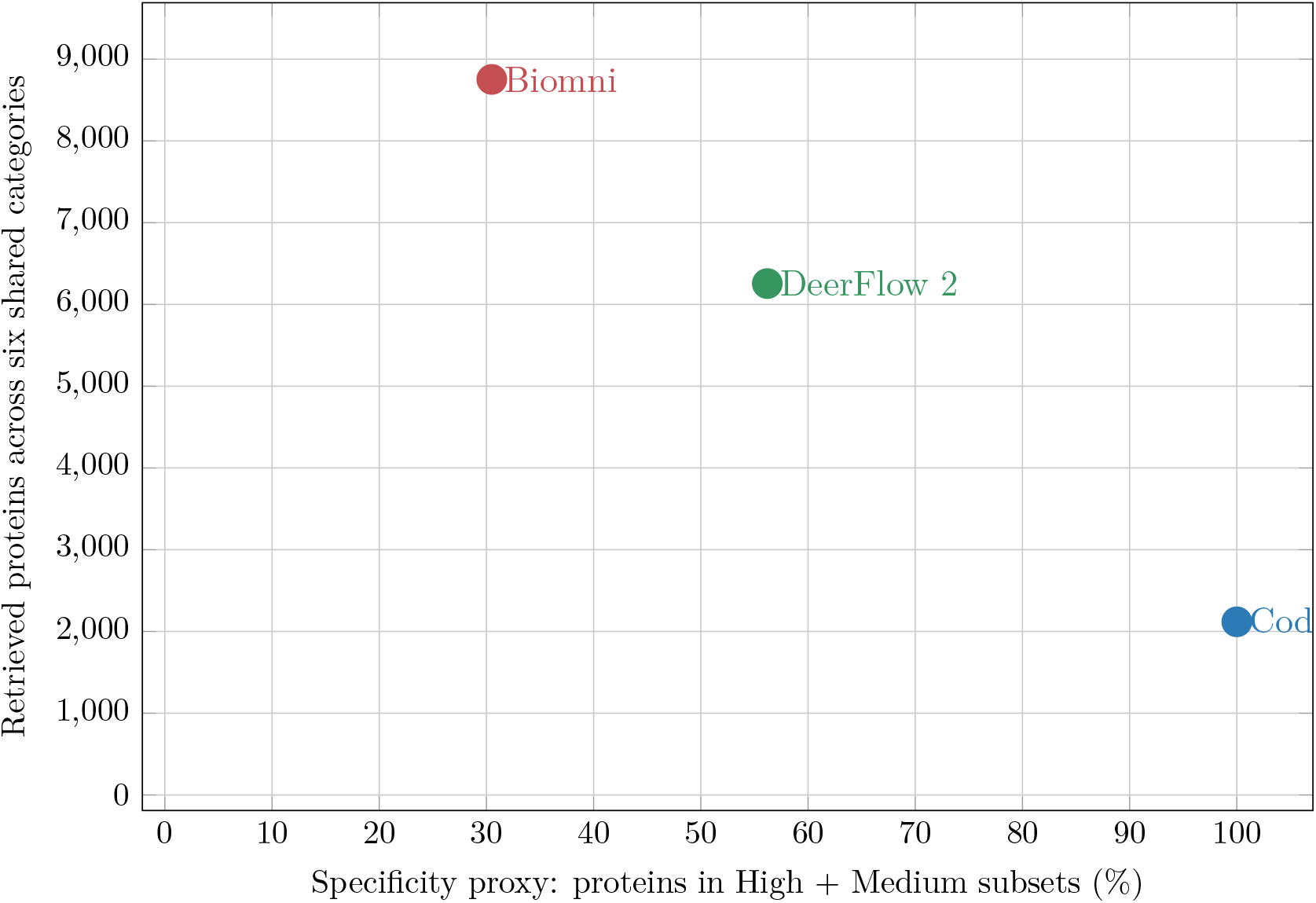
Overall retrieval trade-off across the six shared categories. Higher y-values indicate broader sensitivity; farther right indicates tighter alignment to the benchmark prompt under the subset relevance heuristic. Codex delivered the best balance, DeerFlow was the broadest useful expansion, and Biomni showed the largest but least specific output.

### 3.2 Codex produced the strongest balance of output quality and provenance

Table 1 summarizes the practical profiles of the three systems as observed from their saved artifacts. Codex was the most provenance-rich output: it paired category FASTA files with a manifest, accession-level evidence table, a retrieval script, and a test file. DeerFlow was the most explicit about query generation, exposing exact UniProt query terms in a structured JSON summary and providing per-category TSV files plus a combined deduplicated union. Biomni provided an execution notebook and a broad aggregate CSV, but in this benchmark its functional expansion was substantially less constrained by the calcification prompt.

**Table 1:** System-level comparison of the three agent outputs as observed from the saved run folders.

| System | Benchmark configuration | Observed output artifacts | Practical profile in this benchmark |
| --- | --- | --- | --- |
| Codex + Claude Scientific Skills | OpenAI Codex app extended with the local Claude Scientific Skills library [8, 9] | Per-category FASTA, manifest, evidence tables, retrieval script, test script, evidence summary | Best overall balance of sensitivity and specificity; strongest provenance for downstream curation; best primary source for the final recommended set |
| DeerFlow 2 | DeerFlow 2 with default skills only [12] | Per-category FASTA and TSV, explicit UniProt query terms in `summary.json`, retrieval script, combined deduplicated union, evidence summary | Most exhaustive controlled expansion; especially useful as a targeted supplement for matrix and glycan categories; broader than needed for direct final use |
| Biomni Lab online | Biomni online output compared retrospectively; public repo and preprint cited for system description [10, 11] | Per-category FASTA, aggregate CSV, narrative evidence document, execution-trace notebook | Broadest retrieval but weakest prompt specificity in this task; more suitable for exploratory recall than direct category-resolved export |

### 3.3 Repeated prompts separated the systems by stability

The repeated-run comparison strongly differentiated the systems by reproducibility (Figure 3; Supplementary Tables S4–S6). Codex was nearly invariant across the six categories, with four categories identical between runs and only 56 run-specific proteins across the full benchmark. Its mean category Jaccard was 0.982 and its micro-Jaccard across all category-accession pairs was 0.974. The only substantive change was a 54-protein expansion in category 5, where run 2 added adaptor-complex trafficking proteins that still scored high relevance under the benchmark heuristic. DeerFlow was intermediate, with mean category Jaccard 0.795 and micro-Jaccard 0.571. It was effectively stable in C1 and C2 (J = 1.000 and 0.956), moderately stable in C3–C5 (0.762–0.847), and much less stable in signaling and gene control (C6, J = 0.389), where run 1 produced a much larger generic kinase-rich expansion than run 2. Biomni was the least stable system, with mean category Jaccard 0.412 and micro-Jaccard 0.319. Its inorganic-carbon set was reasonably reproducible (C1, J = 0.822), but calcium/pH homeostasis and signaling shifted sharply between runs (C2, J = 0.166; C6, J = 0.137), and run 1 was far larger than run 2 in every category except C2. Under the stricter within-system heuristic, Codex’s run-specific proteins all remained in high- or medium-relevance subsets, whereas DeerFlow’s instability was concentrated in broad signaling and trafficking expansions and Biomni’s variable proteins were overwhelmingly low relevance.

**Figure 3:**
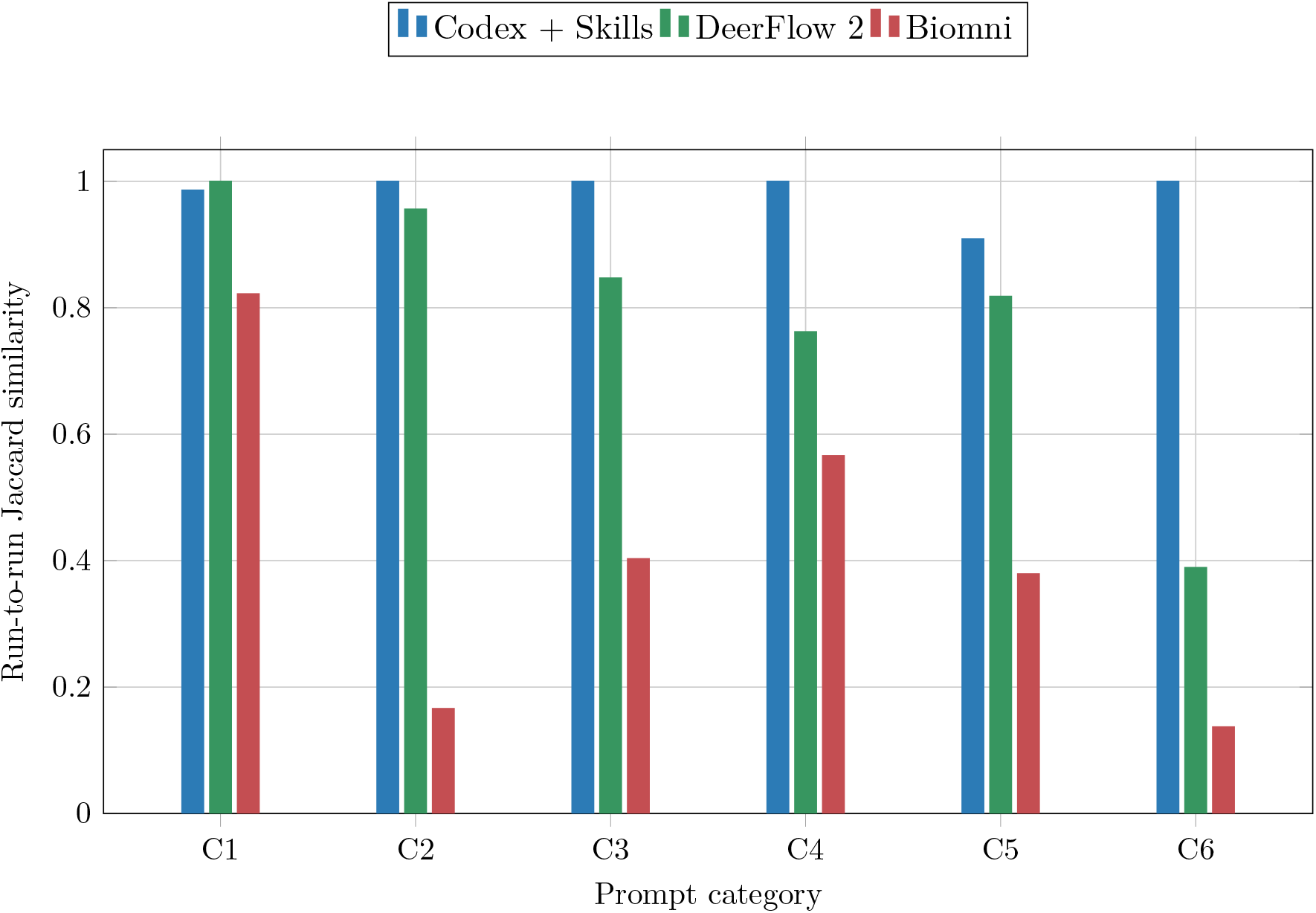
Within-system run-to-run stability across the six shared prompt categories. C1–C6 follow the prompt order from inorganic-carbon acquisition through signaling and gene control. Codex was nearly invariant, DeerFlow was stable in the tighter transport categories but much less stable in signaling, and Biomni showed the largest run-to-run swings in the broadest bins.

### 3.4 Agreement was high only for the most tightly defined category

The category-by-category comparison revealed that agent agreement depended strongly on how tightly the biology matched discrete UniProt annotation families (Figure 4; Table 2). The clearest example was inorganic carbon acquisition, where all three systems converged on a 129-protein triple-overlap core dominated by carbonic anhydrases and bicarbonate transporters. This category had a triple-overlap fraction of 69.0% of the full union, far higher than any other category, indicating that the prompt closely aligned with recognizable UniProt annotations. The complete 42-row overlap decomposition is provided in Supplementary Table S1, the full region-level relevance annotations are provided in Supplementary Tables S2 and S3, and the repeated-run stability summaries are provided in Supplementary Tables S4–S6.

**Table 2:**
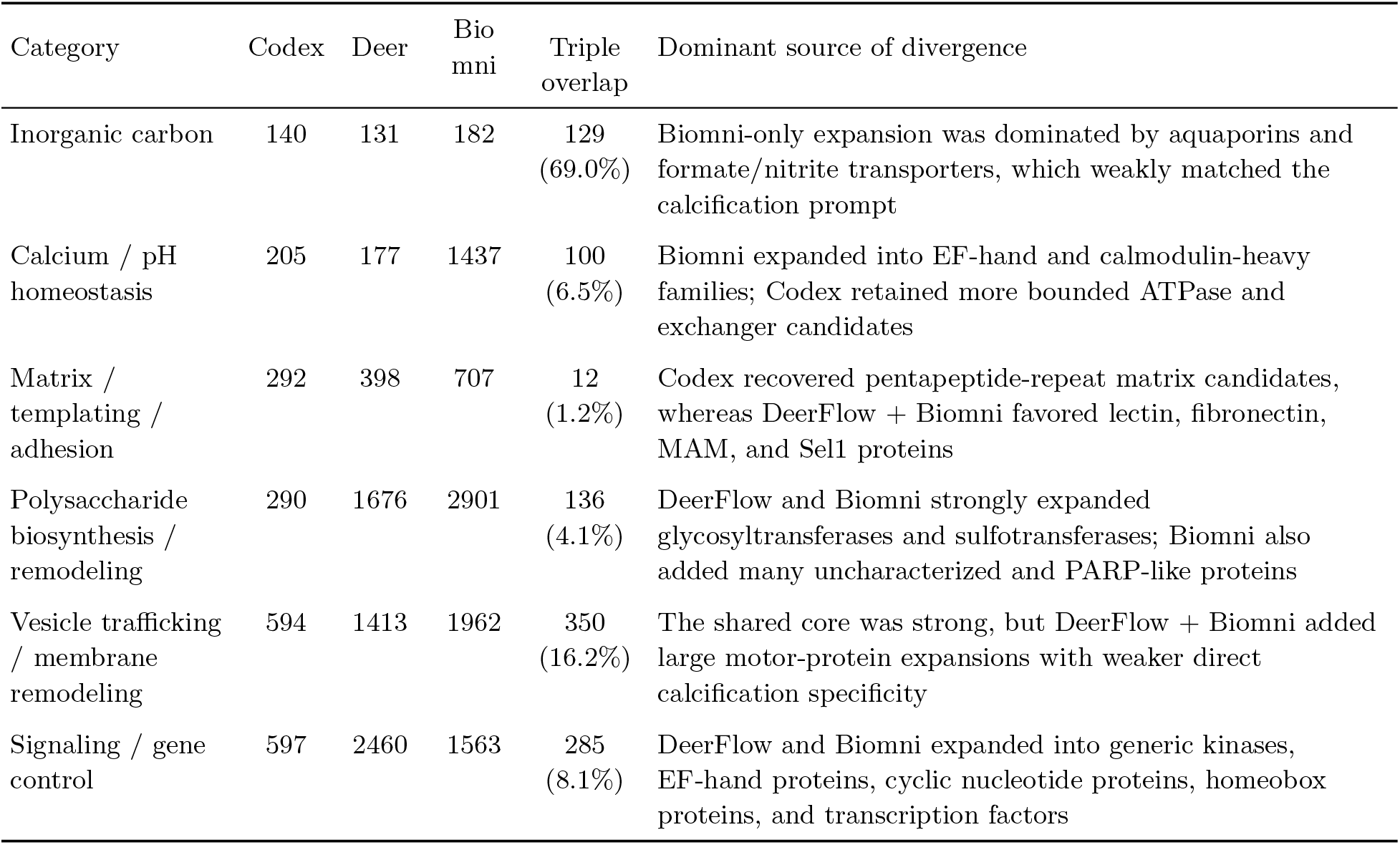
Category-level comparison across the six shared benchmark categories. Triple overlap is reported as a fraction of the union of all three systems for that category.

**Figure 4:**
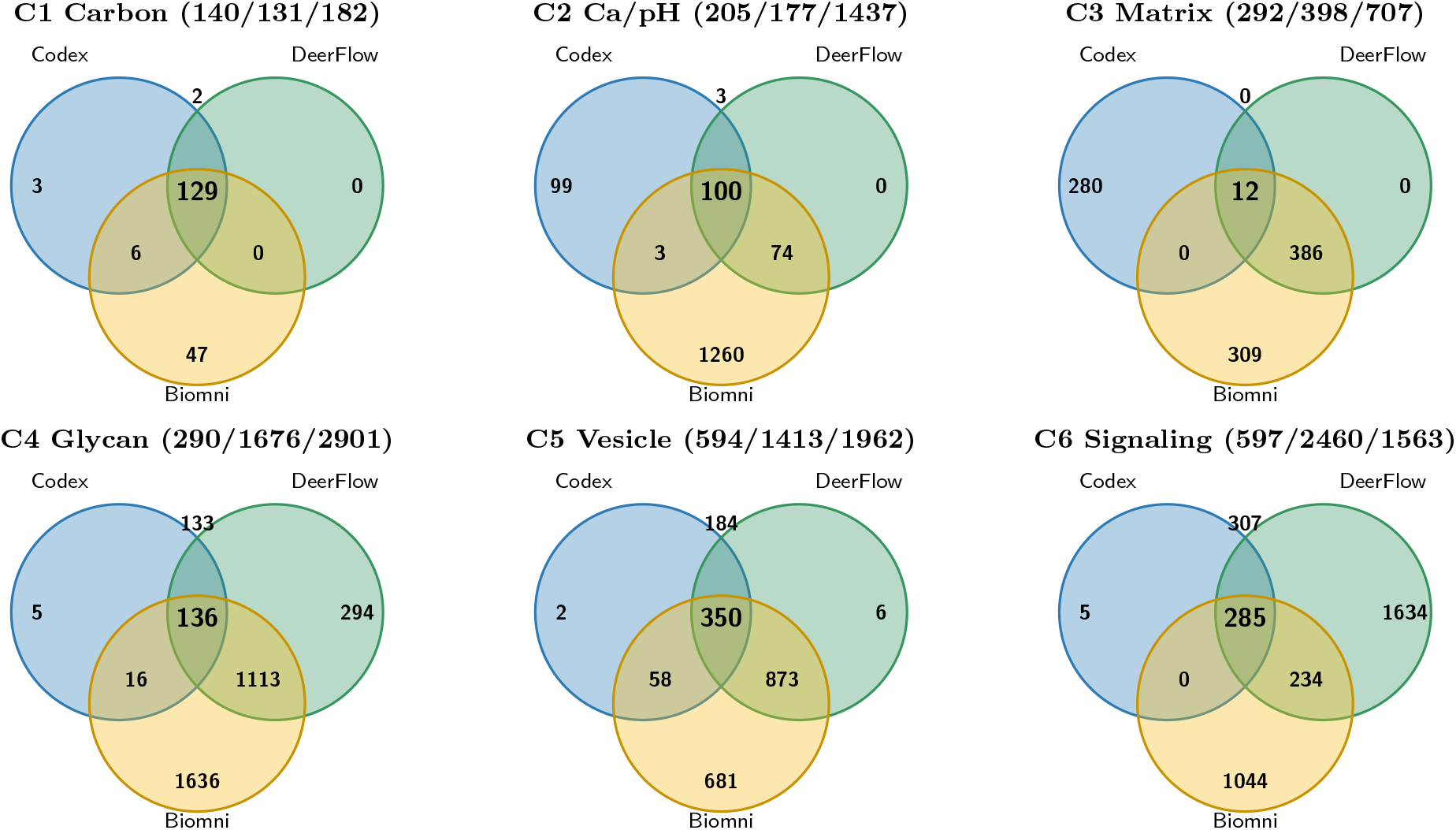
Three-way accession overlap for the six shared benchmark categories in the run 2 cross-agent comparison. Interior numbers report the seven Venn regions for each category, and the panel headers report total proteins retrieved by each agent in that category. Circle areas are schematic and are not proportional to set size.

The remaining categories diverged more sharply. Calcium and pH homeostasis shared a robust 100-protein core dominated by sodium/calcium exchangers and a few calcium-transporting ATPases, but Biomni expanded aggressively into EF-hand proteins and calmodulin-related entries. The organic matrix category showed the strongest conceptual split: Codex recovered 280 pentapeptide-repeat candidates absent from the other outputs, whereas DeerFlow and Biomni shared 386 lectin-, fibronectin-, MAM-, and Sel1-domain proteins that also matched the prompt well. Matrix-polysaccharide remodeling and vesicle trafficking both contained strong shared cores but showed large expansions in DeerFlow and Biomni. Signaling and gene control was the broadest and least specific category overall, confirming the benchmark prompt’s own caution that transcriptional and signaling regulators are harder to retrieve cleanly than transporters or matrix proteins.

### 3.5 Category-specific comparisons clarified how to build a balanced final set

The six categories separated naturally into three benchmarking patterns. First, the tightly defined transport categories favored Codex. In inorganic carbon acquisition, nearly the entire useful core was shared, but the only substantial extra expansion came from Biomni and was mostly weakly aligned. In calcium and proton handling, Codex added 99 unique P-type ATPase and vacuolar ATPase candidates that remained plausible and substantially cleaner than Biomni’s EF-hand expansion. Second, matrix and glycan categories benefited from DeerFlow supplementation. In the organic matrix category, the prompt explicitly allowed inferential inclusion of adhesion-like proteins, and DeerFlow recovered many lectin, fibronectin, MAM, and Sel1 candidates that complemented Codex’s pentapeptide-repeat set. In the glycan-remodeling category, DeerFlow uniquely added 294 glycosyltransferase-rich proteins that were still highly relevant, whereas Biomni’s much larger additions contained a heavier burden of uncharacterized and weakly connected proteins. Third, signaling should be treated as a conservative bin. Although all three systems shared a 285-protein core dominated by calcineurin-like proteins, 14-3-3 proteins, phospholipase C, and calcium-dependent protein kinases, the larger DeerFlow and Biomni expansions rapidly drifted toward generic signaling machinery.

These results directly motivated the balanced recommendation generated from the benchmark. The best final set used Codex as the backbone for categories 1, 2, 4, 5, and 6, while integrating DeerFlow selectively into category 3 for its adhesion- and templating-domain candidates. In other words, the best practical answer was not the output of any one system exactly as produced, but a hybrid assembled from the most biologically coherent regions of the overlap analysis.

## 4 Discussion

This benchmark suggests that for large protein retrieval tasks with complex functional structure, agent performance depends more on workflow design than on raw output volume. Codex was not the most exhaustive system, yet it was the most useful because its retrievals remained close to the benchmark’s biological intent. DeerFlow was the most valuable expansion engine because it made its query structure explicit and tended to broaden within still-relevant family space, especially for matrix and carbohydrate biology. Biomni, despite being a biomedical specialist agent, behaved in this task more like a high-recall exploratory retriever than a precise protein-set constructor. That result is not a general indictment of Biomni; rather, it reflects a mismatch between a broad biomedical reasoning system and a niche task that rewards tightly bounded taxonomic and annotation-aware query generation.

The repeated-run analysis strengthens this interpretation. Codex’s near-deterministic behavior suggests that its bounded retrieval logic and provenance-rich exports are robust to modest prompt-level variation. DeerFlow sits in the middle: when the biology maps to tighter transporter or adhesion families, its outputs are highly reproducible, but when the search space expands into underconstrained signaling or glycan-remodeling space, the same system becomes much more prompt-sensitive. Biomni’s low repeatability is consistent with an exploratory high-recall strategy that can swing widely when broad biomedical families dominate the query. In practice, repeated-run stability is itself a useful benchmark dimension, because unstable broad outputs impose much larger downstream curation costs even when their single best run contains many plausible proteins.

The benchmark also highlights a biological point. Categories tied to well-defined transport or enzyme families are much easier for AI agents to retrieve consistently than categories centered on matrix structure or upstream regulation. Carbonic anhydrases, bicarbonate transporters, sodium/calcium exchangers, V-type ATPases, SNAREs, clathrin-associated factors, and phospholipase C are closer to classical database entities. By contrast, coccolith matrix proteins, adhesive scaffolds, CAP-associated machinery, and developmental regulators remain partly inferential even in the literature [5, 6]. Any agent that tries to be exhaustive in those spaces will inevitably risk either missing plausible candidates or admitting too many generic proteins.

### 4.1 Best-practice recommendations for similar AI bioinformatics tasks

#### Decompose the biology before querying

The six-category prompt worked because it decomposed calcification into concrete subsystems. Large retrieval tasks should be phrased as mechanistic bins rather than one monolithic natural-language request.

#### Force taxonomic scope explicitly

All three systems benefited from coccolithophore-only framing, but differences in taxon scope likely still influenced output size. Prompts of this kind should name target taxa and, when relevant, explicitly exclude known non-target relatives.

#### Request provenance artifacts, not just FASTA files

The most useful output was not simply the most accurate FASTA set; it was the set accompanied by scripts, query terms, manifests, and evidence tables. These artifacts make downstream manual curation far easier.

#### Use a two-stage workflow when sensitivity matters

A practical strategy is to run one tighter agent for the primary set and one broader agent for candidate supplementation, then compare the overlap regions rather than accepting the largest output wholesale.

#### Treat signaling and transcription bins conservatively

Broad regulator categories are highly susceptible to drift into generic kinases, calcium sensors, and transcription factors. For these bins, specificity should be prioritized over recall unless there is strong literature support.

#### Benchmark at the accession level

AI-generated retrievals can sound convincing while still differing greatly in protein identity. Accession-level overlap, not narrative confidence, should be the default comparison unit.

#### Benchmark repeated runs, not just one prompt

The repeated-run analysis here showed that broad categories such as signaling and calcium-regulatory proteins can vary dramatically even within the same system. At least two independent runs should be compared before treating any large AI-retrieved protein set as stable.

### 4.2 Limitations

This study has several limitations. First, it now includes two runs per system, but that is still too small to estimate full within-system variance across broader prompt perturbations, user interventions, or future model-version changes. Second, the comparison focuses on output quality because runtime, token usage, and human interaction burden were not uniformly recorded. Third, the relevance labels are heuristic and literature-informed rather than manually curated accession by accession. Fourth, UniProt annotations in coccolithophores remain incomplete and many proteins are TrEMBL entries, so any system operating purely through annotation matching will inherit those biases [1]. Finally, the benchmark evaluates a very specific class of task: large protein-set retrieval with complex functional annotation. It does not test the same systems on structural biology, transcriptomics, or wet-lab planning tasks.

## 5 Conclusion

In this coccolithophore calcification benchmark, Codex app agents with Claude Scientific Skills produced the best balanced retrieval, DeerFlow 2 provided the most useful targeted expansion, and Biomni Lab online produced the largest but least specific sets. The repeated-run analysis reinforced this ranking: Codex was also the most reproducible, DeerFlow was intermediate, and Biomni was the least stable. The most important lesson is that agent choice alone is not enough. High-quality bioinformatics retrieval depends on mechanistically structured prompts, explicit taxonomic boundaries, provenance-rich outputs, accession-level post hoc comparison, and repeated-run stability checks. For similar tasks, the best practice is a hybrid workflow: use a specificity-oriented agent to establish the backbone, a broader agent to search for category-specific additions, and a transparent overlap analysis to decide what to keep.

## Supplementary Materials

Supplementary tables derived directly from the overlap analysis and repeated-run comparison are provided in the companion supplement supplement.pdf. These include the full overlap decomposition across the six shared categories, the full relevance summary of each Venn subset, dominant protein-family summaries and interpretive notes, category-level repeated-run stability summaries, system-level repeatability metrics, and run-specific subset relevance summaries.

## Data and Code Availability

All benchmark outputs used in this study are available at github.com/xiaoyu12/calc_protein_retrieval.

## Acknowledgments

I thank the developers of Codex, Claude Scientific Skills, Biomni, DeerFlow, and UniProt, as well as the coccolithophore research community whose primary studies made this benchmark biologically interpretable.

## Author Contributions

X.Z. conceived the benchmark, analyzed the outputs, wrote the manuscript, and prepared the comparison artifacts.

## References

[1] The UniProt Consortium. UniProt: the universal protein knowledgebase in 2023. Nucleic Acids Research, 51(D1):D523–D531, 2023. doi: 10.1093/nar/gkac1052. URL https://doi.org/10.1093/nar/gkac1052.

[2] Luke Mackinder, Glen Wheeler, Declan Schroeder, Ulf Riebesell, and Colin Brownlee. Molecular mechanisms underlying calcification in coccolithophores. Geomicrobiology Journal, 27 (6-7):585–595, 2010. doi: 10.1080/01490451003703014. URL https://doi.org/10.1080/01490451003703014.

[3] L. C. M. Mackinder, G. Wheeler, D. Schroeder, P. von Dassow, U. Riebesell, and C. Brownlee. Expression of biomineralization-related ion transport genes in Emiliania huxleyi. Environmental Microbiology, 13(12):3250–3265, 2011. doi: 10.1111/j.1462-2920.2011.02561.x. URL https://doi.org/10.1111/j.1462-2920.2011.02561.x.

[4] Alison R. Taylor, Abdul Chrachri, Glen Wheeler, Helen Goddard, and Colin Brownlee. A voltage-gated H^+^channel underlying ph homeostasis in calcifying coccolithophores. PLoS Biology, 9(6):e1001085, 2011. doi: 10.1371/journal.pbio.1001085. URL https://doi.org/10.1371/journal.pbio.1001085.

[5] Alastair Skeffington, Axel Fischer, Sanja Sviben, et al. A joint proteomic and genomic investigation provides insights into the mechanism of calcification in coccolithophores. Nature Communications, 14:3749, 2023. doi: 10.1038/s41467-023-39336-1. URL https://doi.org/10.1038/s41467-023-39336-1.

[6] Craig J. Dedman, Nishant Chauhan, Alba González-Lanchas, et al. Exploring proteins within the coccolith matrix. Scientific Reports, 14:31821, 2024. doi: 10.1038/s41598-024-83052-9. URL https://doi.org/10.1038/s41598-024-83052-9.

[7] Peter Quinn, R. M. Bowers, Xiaoyan Zhang, et al. cdna microarrays as a tool for identification of biomineralization proteins in the coccolithophorid Emiliania huxleyi (Haptophyta). Applied and Environmental Microbiology, 72(8):5512–5526, 2006. doi: 10.1128/AEM.00343-06. URL https://doi.org/10.1128/AEM.00343-06.

[8] OpenAI. Introducing the Codex app. https://openai.com/index/introducing-the-codex-app/, 2026. Accessed 2026-03-28.

[9] K-Dense-AI. Claude scientific skills. https://github.com/K-Dense-AI/claude-scientific-skills, 2026. Accessed 2026-03-28.

[10] Kexin Huang, Serena Zhang, Hanchen Wang, Yuanhao Qu, et al. Biomni: A general-purpose biomedical AI agent. bioRxiv, 2025. doi: 10.1101/2025.05.30.656746. URL https://www.biorxiv.org/content/10.1101/2025.05.30.656746v1.

[11] SNAP Stanford. Biomni. https://github.com/snap-stanford/Biomni, 2026. Accessed 2026-03-28.

[12] ByteDance. Deerflow 2.0. https://github.com/bytedance/deer-flow, 2026. Accessed 2026-03-28.

[13] Osung Nam, Ichiro Suzuki, Yoshihiro Shiraiwa, and EonSeon Jin. Association of phosphatidylinositol-specific phospholipase C with calcium-induced biomineralization in the coccolithophore Emiliania huxleyi. Microorganisms, 8(9):1389, 2020. doi: 10.3390/microorganisms8091389. URL https://doi.org/10.3390/microorganisms8091389.

[14] Kana Kayano and Yoshihiro Shiraiwa. Physiological regulation of coccolith polysaccharide production by phosphate availability in the coccolithophorid Emiliania huxleyi. Plant and Cell Physiology, 50(8):1522–1531, 2009. doi: 10.1093/pcp/pcp097. URL https://doi.org/10.1093/pcp/pcp097.

