## Supplementary Tables for "Benchmarking Agentic Bioinformatics Systems for Complex Protein-Set Retrieval: A Coccolithophore Calcification Case Study"

This supplement provides table-level outputs derived directly from the accession-level comparison archived in `comparison_results/run_2_detailed_comparison` and the repeated-run variance analysis archived in `comparison_results/repeated_run_variance`. Table S1 reports the full overlap decomposition across all six shared categories. Table S2 reports the full relevance summary for every Venn subset. Table S3 reports the dominant protein families and interpretive notes used to evaluate whether each subset represented biologically coherent recovery or query broadening. Table S4 reports category-level repeated-run stability for each agent. Table S5 reports system-level repeated-run stability metrics. Table S6 reports the relevance of run-specific subsets that appeared in only one of the two repeated runs.

Table 1: Full accession-level overlap decomposition across the six shared benchmark categories. Counts are reported for all seven Venn regions per category after normalization to UniProt accessions.

| Category | Region | Count | Codex | DeerFlow | Biomni |
|---|---|---|---|---|---|
| Inorganic-carbon acquisition and carbonate chemistry | All three | 129 | 140 | 131 | 182 |
| Inorganic-carbon acquisition and carbonate chemistry | Biomni only | 47 | 140 | 131 | 182 |
| Inorganic-carbon acquisition and carbonate chemistry | Codex + Biomni only | 6 | 140 | 131 | 182 |
| Inorganic-carbon acquisition and carbonate chemistry | Codex + deer_flow only | 2 | 140 | 131 | 182 |
| Inorganic-carbon acquisition and carbonate chemistry | Codex only | 3 | 140 | 131 | 182 |
| Inorganic-carbon acquisition and carbonate chemistry | deer_flow + Biomni only | 0 | 140 | 131 | 182 |
| Inorganic-carbon acquisition and carbonate chemistry | deer_flow only | 0 | 140 | 131 | 182 |
| Calcium delivery and proton or pH homeostasis | All three | 100 | 205 | 177 | 1437 |
| Calcium delivery and proton or pH homeostasis | Biomni only | 1260 | 205 | 177 | 1437 |
| Calcium delivery and proton or pH homeostasis | Codex + Biomni only | 3 | 205 | 177 | 1437 |
| Calcium delivery and proton or pH homeostasis | Codex + deer_flow only | 3 | 205 | 177 | 1437 |
| Calcium delivery and proton or pH homeostasis | Codex only | 99 | 205 | 177 | 1437 |
| Calcium delivery and proton or pH homeostasis | deer_flow + Biomni only | 74 | 205 | 177 | 1437 |
| Calcium delivery and proton or pH homeostasis | deer_flow only | 0 | 205 | 177 | 1437 |
| Organic matrix, crystal templating, and adhesion proteins | All three | 12 | 292 | 398 | 707 |
| Organic matrix, crystal templating, and adhesion proteins | Biomni only | 309 | 292 | 398 | 707 |
| Organic matrix, crystal templating, and adhesion proteins | Codex + Biomni only | 0 | 292 | 398 | 707 |
| Organic matrix, crystal templating, and adhesion proteins | Codex + deer_flow only | 0 | 292 | 398 | 707 |
| Organic matrix, crystal templating, and adhesion proteins | Codex only | 280 | 292 | 398 | 707 |
| Organic matrix, crystal templating, and adhesion proteins | deer_flow + Biomni only | 386 | 292 | 398 | 707 |

*Continued on next page*

*Table continued from previous page*

| Category | Region | Count | Codex | DeerFlow | Biomni |
|---|---|---|---|---|---|
| Organic matrix, crystal templating, and adhesion proteins | deer_flow only | 0 | 292 | 398 | 707 |
| Matrix-polysaccharide biosynthesis and remodeling | All three | 136 | 290 | 1676 | 2901 |
| Matrix-polysaccharide biosynthesis and remodeling | Biomni only | 1636 | 290 | 1676 | 2901 |
| Matrix-polysaccharide biosynthesis and remodeling | Codex + Biomni only | 16 | 290 | 1676 | 2901 |
| Matrix-polysaccharide biosynthesis and remodeling | Codex + deer_flow only | 133 | 290 | 1676 | 2901 |
| Matrix-polysaccharide biosynthesis and remodeling | Codex only | 5 | 290 | 1676 | 2901 |
| Matrix-polysaccharide biosynthesis and remodeling | deer_flow + Biomni only | 1113 | 290 | 1676 | 2901 |
| Matrix-polysaccharide biosynthesis and remodeling | deer_flow only | 294 | 290 | 1676 | 2901 |
| Coccolith-vesicle biogenesis, trafficking, and membrane remodeling | All three | 350 | 594 | 1413 | 1962 |
| Coccolith-vesicle biogenesis, trafficking, and membrane remodeling | Biomni only | 681 | 594 | 1413 | 1962 |
| Coccolith-vesicle biogenesis, trafficking, and membrane remodeling | Codex + Biomni only | 58 | 594 | 1413 | 1962 |
| Coccolith-vesicle biogenesis, trafficking, and membrane remodeling | Codex + deer_flow only | 184 | 594 | 1413 | 1962 |
| Coccolith-vesicle biogenesis, trafficking, and membrane remodeling | Codex only | 2 | 594 | 1413 | 1962 |
| Coccolith-vesicle biogenesis, trafficking, and membrane remodeling | deer_flow + Biomni only | 873 | 594 | 1413 | 1962 |
| Coccolith-vesicle biogenesis, trafficking, and membrane remodeling | deer_flow only | 6 | 594 | 1413 | 1962 |
| Signaling and gene-control regulators | All three | 285 | 597 | 2460 | 1563 |
| Signaling and gene-control regulators | Biomni only | 1044 | 597 | 2460 | 1563 |
| Signaling and gene-control regulators | Codex + Biomni only | 0 | 597 | 2460 | 1563 |
| Signaling and gene-control regulators | Codex + deer_flow only | 307 | 597 | 2460 | 1563 |
| Signaling and gene-control regulators | Codex only | 5 | 597 | 2460 | 1563 |
| Signaling and gene-control regulators | deer_flow + Biomni only | 234 | 597 | 2460 | 1563 |
| Signaling and gene-control regulators | deer_flow only | 1634 | 597 | 2460 | 1563 |

Table 2: Full relevance summary for all overlap regions. Positive and caution fractions were computed from prompt-matching and cautionary keyword enrichment, respectively, and then converted to percentages for display.

| Category | Region | Count | Relevance | Positive | Caution |
|---|---|---|---|---|---|
| Inorganic-carbon acquisition and carbonate chemistry | All three | 129 | High | 100.0% | 0.0% |
| Inorganic-carbon acquisition and carbonate chemistry | Biomni only | 47 | Low | 0.0% | 100.0% |

*Continued on next page*

*Table continued from previous page*

| Category | Region | Count | Relevance | Positive | Caution |
|---|---|---|---|---|---|
| Inorganic-carbon acquisition and carbonate chemistry | Codex + Biomni only | 6 | High | 100.0% | 0.0% |
| Inorganic-carbon acquisition and carbonate chemistry | Codex + deer_flow only | 2 | High | 100.0% | 0.0% |
| Inorganic-carbon acquisition and carbonate chemistry | Codex only | 3 | Medium | 66.7% | 0.0% |
| Inorganic-carbon acquisition and carbonate chemistry | deer_flow + Biomni only | 0 | Low | 0.0% | 0.0% |
| Inorganic-carbon acquisition and carbonate chemistry | deer_flow only | 0 | Low | 0.0% | 0.0% |
| Calcium delivery and proton or pH homeostasis | All three | 100 | High | 100.0% | 0.0% |
| Calcium delivery and proton or pH homeostasis | Biomni only | 1260 | Low | 10.8% | 84.6% |
| Calcium delivery and proton or pH homeostasis | Codex + Biomni only | 3 | High | 100.0% | 0.0% |
| Calcium delivery and proton or pH homeostasis | Codex + deer_flow only | 3 | High | 100.0% | 0.0% |
| Calcium delivery and proton or pH homeostasis | Codex only | 99 | Medium | 100.0% | 0.0% |
| Calcium delivery and proton or pH homeostasis | deer_flow + Biomni only | 74 | High | 100.0% | 0.0% |
| Calcium delivery and proton or pH homeostasis | deer_flow only | 0 | Low | 0.0% | 0.0% |
| Organic matrix, crystal templating, and adhesion proteins | All three | 12 | High | 100.0% | 0.0% |
| Organic matrix, crystal templating, and adhesion proteins | Biomni only | 309 | Low | 1.0% | 95.8% |
| Organic matrix, crystal templating, and adhesion proteins | Codex + Biomni only | 0 | Low | 0.0% | 0.0% |
| Organic matrix, crystal templating, and adhesion proteins | Codex + deer_flow only | 0 | Low | 0.0% | 0.0% |
| Organic matrix, crystal templating, and adhesion proteins | Codex only | 280 | High | 100.0% | 0.0% |
| Organic matrix, crystal templating, and adhesion proteins | deer_flow + Biomni only | 386 | High | 100.0% | 0.0% |
| Organic matrix, crystal templating, and adhesion proteins | deer_flow only | 0 | Low | 0.0% | 0.0% |
| Matrix-polysaccharide biosynthesis and remodeling | All three | 136 | High | 100.0% | 0.0% |
| Matrix-polysaccharide biosynthesis and remodeling | Biomni only | 1636 | Low | 21.2% | 16.3% |
| Matrix-polysaccharide biosynthesis and remodeling | Codex + Biomni only | 16 | High | 100.0% | 0.0% |
| Matrix-polysaccharide biosynthesis and remodeling | Codex + deer_flow only | 133 | High | 100.0% | 0.0% |
| Matrix-polysaccharide biosynthesis and remodeling | Codex only | 5 | High | 80.0% | 0.0% |
| Matrix-polysaccharide biosynthesis and remodeling | deer_flow + Biomni only | 1113 | High | 99.2% | 4.0% |
| Matrix-polysaccharide biosynthesis and remodeling | deer_flow only | 294 | High | 100.0% | 0.0% |
| Coccolith-vesicle biogenesis, trafficking, and membrane remodeling | All three | 350 | High | 100.0% | 0.0% |
| Coccolith-vesicle biogenesis, trafficking, and membrane remodeling | Biomni only | 681 | Low | 5.9% | 64.5% |
| Coccolith-vesicle biogenesis, trafficking, and membrane remodeling | Codex + Biomni only | 58 | Medium | 100.0% | 0.0% |

*Continued on next page*

*Table continued from previous page*

| Category | Region | Count | Relevance | Positive | Caution |
|---|---|---|---|---|---|
| Coccolith-vesicle biogenesis, trafficking, and membrane remodeling | Codex + deer_flow only | 184 | High | 100.0% | 0.0% |
| Coccolith-vesicle biogenesis, trafficking, and membrane remodeling | Codex only | 2 | High | 100.0% | 0.0% |
| Coccolith-vesicle biogenesis, trafficking, and membrane remodeling | deer_flow + Biomni only | 873 | Low | 28.8% | 71.2% |
| Coccolith-vesicle biogenesis, trafficking, and membrane remodeling | deer_flow only | 6 | High | 100.0% | 0.0% |
| Signaling and gene-control regulators | All three | 285 | High | 100.0% | 0.0% |
| Signaling and gene-control regulators | Biomni only | 1044 | Low | 20.9% | 83.8% |
| Signaling and gene-control regulators | Codex + Biomni only | 0 | Low | 0.0% | 0.0% |
| Signaling and gene-control regulators | Codex + deer_flow only | 307 | High | 100.0% | 0.0% |
| Signaling and gene-control regulators | Codex only | 5 | High | 100.0% | 0.0% |
| Signaling and gene-control regulators | deer_flow + Biomni only | 234 | Low-Medium | 67.9% | 32.1% |
| Signaling and gene-control regulators | deer_flow only | 1634 | Low-Medium | 99.4% | 0.0% |

Table 3: Dominant protein families and interpretive notes for each overlap region. These notes were used to distinguish biologically coherent expansions from query broadening.

| Category | Region | Top proteins | Interpretation |
|---|---|---|---|
| Inorganic-carbon acquisition and carbonate chemistry | All three | Bicarbonate transporter-like transmembrane domain-containing protein (48); Carbonic anhydrase (EC 4.2.1.1) (42); Carbonic anhydrase (EC 4.2.1.1) (Carbonate dehydratase) (20); Carbonic anhydrase (9); Gamma carbonic anhydrase (4); Gamma carbonic anhydrase family protein (3) | cross-agent core; strongly enriched for prompt-matching families; dominant positive terms: carbonic anhydrase (81), bicarbonate (48) |
| Inorganic-carbon acquisition and carbonate chemistry | Biomni only | Aquaporin (32); Formate/nitrite transporter (6); Putative formate/nitrite transporter (3); Major intrinsic protein (2); Sulfate transporter (2); Formate/nitrite transporter family protein (1) | Biomni-unique subset; is weakly aligned to the prompt; caution terms: aquaporin (32), formate (11), nitrite (11), major intrinsic protein (2); this looks like query broadening rather than a calcification-focused expansion |
| Inorganic-carbon acquisition and carbonate chemistry | Codex + Biomni only | Carbonic anhydrase (EC 4.2.1.1) (Carbonate dehydratase) (3); carbonic anhydrase (EC 4.2.1.1) (2); Putative gamma-type carbonic anhydrase (1) | paired support without deer_flow; strongly enriched for prompt-matching families; dominant positive terms: carbonic anhydrase (6) |
| Inorganic-carbon acquisition and carbonate chemistry | Codex + deer_flow only | HCO3 transporter (2) | paired support without Biomni; strongly enriched for prompt-matching families; dominant positive terms: hco3 (2) |
| Inorganic-carbon acquisition and carbonate chemistry | Codex only | Carbonate dehydratase-like protein (1); HCO3–transporter (1); Tentative Ca2+ interacting HCO3-transporter (1) | Codex-unique subset; contains many prompt-matching families but is less clean; dominant positive terms: hco3 (2) |

*Continued on next page*

*Table continued from previous page*

| Category | Region | Top proteins | Interpretation |
|---|---|---|---|
| Inorganic-carbon acquisition and carbonate chemistry | deer_flow + Biomni only | | No proteins fall into this region. |
| Inorganic-carbon acquisition and carbonate chemistry | deer_flow only | | No proteins fall into this region. |
| Calcium delivery and proton or pH homeostasis | All three | Sodium/calcium exchanger membrane region domain-containing protein (97); Calcium-transporting ATPase (EC 7.2.2.10) (2); Sodium/Calcium exchanger protein (1) | cross-agent core; strongly enriched for prompt-matching families; dominant positive terms: calcium (100), sodium/calcium (98), calcium-transporting atpase (2) |
| Calcium delivery and proton or pH homeostasis | Biomni only | EF-hand domain-containing protein (804); Calmodulin-lysine N-methyltransferase (80); Cation/H+ exchanger transmembrane domain-containing protein (75); Calmodulin (62); Cation/H+ exchanger domain-containing protein (51); Ion transport domain-containing protein (17) | Biomni-unique subset; is weakly aligned to the prompt; dominant positive terms: cation/h+ (126), calcium (6), hydrogen channel (4); caution terms: ef-hand (808), calmodulin (143), calmodulin-lysine (80), atp synthase (54); this looks like query broadening rather than a calcification-focused expansion |
| Calcium delivery and proton or pH homeostasis | Codex + Biomni only | Cation-transporting P-type ATPase C-terminal domain-containing protein (1); Vacuolar H+-ATPase V1 sector, subunit F (1); Voltage-gated hydrogen channel 1 (Hydrogen voltage-gated channel 1) (1) | paired support without deer_flow; strongly enriched for prompt-matching families; dominant positive terms: cation-transporting p-type atpase (1), hydrogen channel (1), p-type atpase (1), vacuolar h+ (1) |
| Calcium delivery and proton or pH homeostasis | Codex + deer_flow only | Vacuolar ATPase assembly protein VMA22 (3) | paired support without Biomni; strongly enriched for prompt-matching families; dominant positive terms: vma22 (3) |
| Calcium delivery and proton or pH homeostasis | Codex only | Cation-transporting P-type ATPase C-terminal domain-containing protein (28); Cation-transporting P-type ATPase N-terminal domain-containing protein (27); P-type ATPase A domain-containing protein (20); P-type ATPase C-terminal domain-containing protein (11); P-type ATPase N-terminal domain-containing protein (8); Vacuolar H+-ATPase V0 sector, subunits c (2) | Codex-unique subset; strongly enriched for prompt-matching families; dominant positive terms: p-type atpase (94), cation-transporting p-type atpase (55), vacuolar h+ (5); these are mostly ATPase-domain calcium or proton transport candidates that fit the prompt but were not recovered by the broader runs |
| Calcium delivery and proton or pH homeostasis | deer_flow + Biomni only | V-type proton ATPase subunit a (20); V-type proton ATPase proteolipid subunit (17); V-type proton ATPase subunit G (8); V-type proton ATPase subunit S1/VOA1 transmembrane domain-containing protein (8); V-type proton ATPase subunit (6); V-type proton ATPase subunit C (5) | paired expansion without Codex; strongly enriched for prompt-matching families; dominant positive terms: v-type proton atpase (74) |
| Calcium delivery and proton or pH homeostasis | deer_flow only | | No proteins fall into this region. |

*Continued on next page*

*Table continued from previous page*

| Category | Region | Top proteins | Interpretation |
|---|---|---|---|
| Organic matrix, crystal templating, and adhesion proteins | All three | C-type lectin domain-containing protein (4); Fibronectin type-II domain-containing protein (3); Putative calcium binding protein (2); Fibronectin type III-like domain-containing protein (1); Polysacharide-associated, calcium-binding protein (Putative calcium binding protein) (1); Ricin B lectin domain-containing protein (1) | cross-agent core; strongly enriched for prompt-matching families; dominant positive terms: lectin (5), fibronectin (4), calcium binding (3), polysacharide-associated (1) |
| Organic matrix, crystal templating, and adhesion proteins | Biomni only | EGF-like domain-containing protein (170); Tetratricopeptide repeat protein (102); Glutamate-rich WD repeat-containing protein 1 (7); Tetratricopeptide repeat protein 38 (5); Galectin (3); Laminin EGF-like domain-containing protein (3) | Biomni-unique subset; is weakly aligned to the prompt; dominant positive terms: lectin (3); caution terms: egf-like (173), tetratricopeptide (121), von willebrand (2); this looks like query broadening rather than a calcification-focused expansion |
| Organic matrix, crystal templating, and adhesion proteins | Codex + Biomni only | | No proteins fall into this region. |
| Organic matrix, crystal templating, and adhesion proteins | Codex + deer_flow only | | No proteins fall into this region. |
| Organic matrix, crystal templating, and adhesion proteins | Codex only | Pentapeptide repeat-containing protein (264); Pentapeptide repeat protein (16) | Codex-unique subset; strongly enriched for prompt-matching families; dominant positive terms: pentapeptide (280); these Codex-specific hits are useful because they capture pentapeptide-repeat matrix candidates absent from the other outputs |
| Organic matrix, crystal templating, and adhesion proteins | deer_flow + Biomni only | C-type lectin domain-containing protein (105); Fibronectin type-III domain-containing protein (102); MAM domain-containing protein (78); Sel1 repeat family protein (28); Fibronectin type III-like domain-containing protein (25); L-type lectin-like domain-containing protein (10) | paired expansion without Codex; strongly enriched for prompt-matching families; dominant positive terms: lectin (141), fibronectin (135), mam (78), sel1 (32) |
| Organic matrix, crystal templating, and adhesion proteins | deer_flow only | | No proteins fall into this region. |
| Matrix-polysaccharide biosynthesis and remodeling | All three | Glycoside hydrolase family 5 domain-containing protein (78); Glycosyltransferase 2-like domain-containing protein (38); Glycosyltransferase, family 8 (7); Glycoside hydrolase family 31 TIM barrel domain-containing protein (5); Glycoside hydrolase family 31 N-terminal domain-containing protein (4); Sulfotransferase domain-containing protein (2) | cross-agent core; strongly enriched for prompt-matching families; dominant positive terms: glycoside hydrolase (88), glycosyltransferase (46), sulfotransferase (2) |

*Continued on next page*

*Table continued from previous page*

| Category | Region | Top proteins | Interpretation |
|---|---|---|---|
| Matrix-polysaccharide biosynthesis and remodeling | Biomni only | Sulfatase N-terminal domain-containing protein (299); Uncharacterized protein (143); Hexosyltransferase (EC 2.4.1.-) (97); Hydroxyproline O-arabinosyltransferase-like domain-containing protein (86); beta-glucosidase (EC 3.2.1.21) (74); Poly [ADP-ribose] polymerase (PARP) (EC 2.4.2.-) (65) | Biomni-unique subset; is weakly aligned to the prompt; dominant positive terms: sulfatase (334), glycosyltransferase (8), glycosyl hydrolase (5); caution terms: uncharacterized (143), parp (70), poly [adp-ribose] (65), adp-ribosyltransferase (54) |
| Matrix-polysaccharide biosynthesis and remodeling | Codex + Biomni only | Sulfatase N-terminal domain-containing protein (16) | paired support without deer_flow; strongly enriched for prompt-matching families; dominant positive terms: sulfatase (16) |
| Matrix-polysaccharide biosynthesis and remodeling | Codex + deer_flow only | Glycosyltransferase 2-like domain-containing protein (118); Glycosyltransferase, family 8 (6); Glycosyltransferase (5); Family 2 glycosyl transferase (Glycosyltransferase, family 2) (1); Glycosyltransferase family 8 protein (1); Glycosyltransferase, family 2 (1) | paired support without Biomni; strongly enriched for prompt-matching families; dominant positive terms: glycosyltransferase (133) |
| Matrix-polysaccharide biosynthesis and remodeling | Codex only | Asl1-like glycosyl hydrolase catalytic domain-containing protein (2); Glycosyl hydrolase family 92 domain-containing protein (2); Alpha-L-arabinofuranosidase 1 catalytic domain-containing protein (1) | Codex-unique subset; strongly enriched for prompt-matching families; dominant positive terms: glycosyl hydrolase (4) |
| Matrix-polysaccharide biosynthesis and remodeling | deer_flow + Biomni only | Sulfotransferase domain-containing protein (435); Sulfotransferase (83); O-fucosyltransferase family protein (54); Protein-tyrosine sulfotransferase (44); Glycoside hydrolase family 2 catalytic domain-containing protein (41); Glycoside hydrolase family 5 domain-containing protein (41) | paired expansion without Codex; strongly enriched for prompt-matching families; dominant positive terms: sulfotransferase (578), glycoside hydrolase (197), fucosyltransferase (93), mannosidase (69); caution terms: protein-tyrosine sulfotransferase (44) |
| Matrix-polysaccharide biosynthesis and remodeling | deer_flow only | Alpha 1,4-glycosyltransferase domain-containing protein (67); Glycosyltransferase family 92 protein (47); Glycosyltransferase 61 catalytic domain-containing protein (43); Glycosyltransferase 2-like domain-containing protein (29); Glycosyltransferase family 28 N-terminal domain-containing protein (19); Glycosyltransferase subfamily 4-like N-terminal domain-containing protein (17) | deer_flow-unique subset; strongly enriched for prompt-matching families; dominant positive terms: glycosyltransferase (271), glucosyltransferase (19), deacetylase (3), polysaccharide (3) |
| Coccolith-vesicle biogenesis, trafficking, and membrane remodeling | All three | t-SNARE coiled-coil homology domain-containing protein (76); V-SNARE coiled-coil homology domain-containing protein (37); Dynamin-type G domain-containing protein (35); Clathrin light chain (21); Vesicle transport v-SNARE N-terminal domain-containing protein (21); Clathrin/coatomer adaptor adaptin-like N-terminal domain-containing protein (16) | cross-agent core; strongly enriched for prompt-matching families; dominant positive terms: snare (138), coatomer (67), dynamin (58), clathrin (57) |

*Continued on next page*

*Table continued from previous page*

| Category | Region | Top proteins | Interpretation |
|---|---|---|---|
| Coccolith-vesicle biogenesis, trafficking, and membrane remodeling | Biomni only | PX domain-containing protein (138); ATP-grasp domain-containing protein (110); Phosphoinositide phospholipase C (EC 3.1.4.11) (109); Calponin-homology (CH) domain-containing protein (34); ADF-H domain-containing protein (26); ADP-ribosylation factor (21) | Biomni-unique subset; is weakly aligned to the prompt; dominant positive terms: sec23 (32), ap-1 (5), rab (3); caution terms: px domain (138), atp-grasp (131), phosphoinositide phospholipase c (110), calponin-homology (34); this looks like query broadening rather than a calcification-focused expansion |
| Coccolith-vesicle biogenesis, trafficking, and membrane remodeling | Codex + Biomni only | Adaptor protein complex 1 or 2 subunit beta (10); Adaptor protein complex 1 subunit gamma (8); Adaptor protein complex 1 subunit mu (5); Adaptor protein complex 2 subunit alpha (5); Adaptor protein complex 4 subunit mu (5); AP-2 complex subunit alpha (4) | paired support without deer_flow; strongly enriched for prompt-matching families; dominant positive terms: adaptor protein complex (54), ap-2 (4); these adaptor-complex proteins are relevant trafficking machinery even though deer_flow did not recover them here |
| Coccolith-vesicle biogenesis, trafficking, and membrane remodeling | Codex + deer_flow only | Rab-GAP TBC domain-containing protein (124); Rab GDP dissociation inhibitor (13); Beta-adaptin appendage C-terminal subdomain domain-containing protein (9); Small COPII coat GTPase SAR1 (8); Alpha adaptin ear domain protein (5); Rab proteins geranylgeranyltransferase component A (4) | paired support without Biomni; strongly enriched for prompt-matching families; dominant positive terms: rab (156), adaptin (20), copi (8), copii (8) |
| Coccolith-vesicle biogenesis, trafficking, and membrane remodeling | Codex only | Putative rabgap/tbc domain-containing protein (1); Vesicle-trafficking protein SEC22b (1) | Codex-unique subset; strongly enriched for prompt-matching families; dominant positive terms: rab (1), vesicle (1) |
| Coccolith-vesicle biogenesis, trafficking, and membrane remodeling | deer_flow + Biomni only | Myosin motor domain-containing protein (237); Kinesin motor domain-containing protein (200); Kinesin light chain (106); Kinesin-like protein (75); Dynein heavy chain (25); Dynein heavy chain C-terminal domain-containing protein (23) | paired expansion without Codex; has mixed relevance with a smaller calcification signal; dominant positive terms: myosin (245), vesicle (6); caution terms: kinesin (390), dynein (232) |
| Coccolith-vesicle biogenesis, trafficking, and membrane remodeling | deer_flow only | Sorting nexin/Vps5-like C-terminal domain-containing protein (5); Sorting nexin 1/Sorting nexin 2/Vacuolar protein sorting protein 5 (1) | deer_flow-unique subset; strongly enriched for prompt-matching families; dominant positive terms: sorting nexin (6) |
| Signaling and gene-control regulators | All three | Calcineurin-like phosphoesterase domain-containing protein (239); 14-3-3 domain-containing protein (22); Phosphatidylinositol-specific phospholipase C X domain-containing protein (7); Phospholipase C (7); Calcium-dependent protein kinase (4); 14-3-3 protein (3) | cross-agent core; strongly enriched for prompt-matching families; dominant positive terms: calcineurin (239), 14-3-3 (25), phospholipase c (14), calcium-dependent protein kinase (4) |
| Signaling and gene-control regulators | Biomni only | Myb-like domain-containing protein (238); Cyclic nucleotide-binding domain-containing protein (229); Homeobox domain-containing protein (116); Histidine kinase/HSP90-like ATPase domain-containing protein (64); histidine kinase (EC 2.7.13.3) (57); Guanylate cyclase domain-containing protein (55) | Biomni-unique subset; is weakly aligned to the prompt; dominant positive terms: kinase (212), receptor (7); caution terms: myb (271), cyclic nucleotide-binding (230), histidine kinase (135), homeobox (116); this looks like query broadening rather than a calcification-focused expansion |

*Continued on next page*

*Table continued from previous page*

| Category | Region | Top proteins | Interpretation |
|---|---|---|---|
| Signaling and gene-control regulators | Codex + Biomni only | | No proteins fall into this region. |
| Signaling and gene-control regulators | Codex + deer_flow only | Phosphoinositide phospholipase C (EC 3.1.4.11) (109); Calmodulin-lysine N-methyltransferase (80); Calmodulin (62); EF-hand domain-containing protein (52); PP2A regulatory subunit B" EF-hand domain-containing protein (2); Calmodulin-like protein (1) | paired support without Biomni; strongly enriched for prompt-matching families; dominant positive terms: calmodulin (143), phosphoinositide phospholipase c (110), phospholipase c (110), ef-hand (54) |
| Signaling and gene-control regulators | Codex only | Serine-threonine kinase receptor-associated protein (5) | Codex-unique subset; strongly enriched for prompt-matching families; dominant positive terms: kinase (5), receptor (5); this supports the earlier decision to prefer Codex for a tighter signaling set |
| Signaling and gene-control regulators | deer_flow + Biomni only | SAM domain-containing protein (157); Transcription factor CBF/NF-Y/archaeal histone domain-containing protein (27); General transcription factor IIH subunit 3 (7); Transcription factor IIIC subunit 5 HTH domain-containing protein (5); General transcription factor IIH subunit (4); General transcription factor IIH subunit 4 (4) | paired expansion without Codex; contains many prompt-matching families but is less clean; dominant positive terms: sam domain (159); caution terms: transcription factor (75), general transcription factor (18) |
| Signaling and gene-control regulators | deer_flow only | EF-hand domain-containing protein (1174); non-specific serine/threonine protein kinase (EC 2.7.11.1) (407); Non-specific serine/threonine protein kinase (15); Serine/threonine-protein kinase (4); Serine/threonine-protein kinase RIO1 (EC 2.7.11.1) (Serine/threonine-protein kinase rio1) (4); Mitochondrial proton/calcium exchanger protein (Leucine zipper-EF-hand-containing transmembrane protein 1) (3) | deer_flow-unique subset; strongly enriched for prompt-matching families; dominant positive terms: ef-hand (1181), kinase (443), serine/threonine (443); despite matching signaling keywords, this region is dominated by very broad EF-hand and nonspecific kinase expansions rather than tightly calcification-focused regulators |

Table 4: Category-level repeated-run stability for each agent, comparing the accession sets exported in run 1 and run 2 after normalization to UniProt accessions.

| Agent | Cat. | Run 1 | Run 2 | Shared | Union | Jaccard | R1 rec. | R2 rec. |
|---|---|---|---|---|---|---|---|---|
| Codex + Skills | C1 | 138 | 140 | 138 | 140 | 0.986 | 1.000 | 0.986 |
| Codex + Skills | C2 | 205 | 205 | 205 | 205 | 1.000 | 1.000 | 1.000 |
| Codex + Skills | C3 | 292 | 292 | 292 | 292 | 1.000 | 1.000 | 1.000 |
| Codex + Skills | C4 | 290 | 290 | 290 | 290 | 1.000 | 1.000 | 1.000 |
| Codex + Skills | C5 | 540 | 594 | 540 | 594 | 0.909 | 1.000 | 0.909 |
| Codex + Skills | C6 | 597 | 597 | 597 | 597 | 1.000 | 1.000 | 1.000 |
| DeerFlow 2 | C1 | 131 | 131 | 131 | 131 | 1.000 | 1.000 | 1.000 |
| DeerFlow 2 | C2 | 179 | 177 | 174 | 182 | 0.956 | 0.972 | 0.983 |
| DeerFlow 2 | C3 | 461 | 398 | 394 | 465 | 0.847 | 0.855 | 0.990 |
| DeerFlow 2 | C4 | 1689 | 1676 | 1455 | 1910 | 0.762 | 0.861 | 0.868 |
| DeerFlow 2 | C5 | 1636 | 1413 | 1372 | 1677 | 0.818 | 0.839 | 0.971 |
| DeerFlow 2 | C6 | 5449 | 2460 | 2214 | 5695 | 0.389 | 0.406 | 0.900 |
| Biomni | C1 | 217 | 182 | 180 | 219 | 0.822 | 0.829 | 0.989 |
| Biomni | C2 | 915 | 1437 | 335 | 2017 | 0.166 | 0.366 | 0.233 |

*Continued on next page*

*Table continued from previous page*

| Agent | Cat. | Run 1 | Run 2 | Shared | Union | Jaccard | R1 rec. | R2 rec. |
|---|---|---|---|---|---|---|---|---|
| Biomni | C3 | 1714 | 707 | 696 | 1725 | 0.403 | 0.406 | 0.984 |
| Biomni | C4 | 4905 | 2901 | 2822 | 4984 | 0.566 | 0.575 | 0.973 |
| Biomni | C5 | 3206 | 1962 | 1419 | 3749 | 0.379 | 0.443 | 0.723 |
| Biomni | C6 | 7241 | 1563 | 1059 | 7745 | 0.137 | 0.146 | 0.678 |

Table 5: System-level repeated-run stability metrics. Mean Jaccard summarizes category-level repeatability, whereas micro-Jaccard summarizes repeatability across all category-accession pairs pooled together.

| Agent | Run 1 | Run 2 | Mean J | Micro J | Variable | High/med. | Most stable | Most variable |
|---|---|---|---|---|---|---|---|---|
| Codex + Skills | 2062 | 2118 | 0.982 | 0.974 | 56 | 100.0% | C2, C3 | C5, C1 |
| DeerFlow 2 | 9545 | 6255 | 0.795 | 0.571 | 4320 | 11.9% | C1, C2 | C6, C4 |
| Biomni | 18198 | 8752 | 0.412 | 0.319 | 13928 | 0.0% | C1, C4 | C6, C2 |

Table 6: Run-specific subset relevance for repeated-run variance. Only accession subsets unique to run 1 or unique to run 2 are shown, because shared subsets represent the stable cross-run core.

| Agent | Cat. | Subset | Count | Relevance | Positive | Caution |
|---|---|---|---|---|---|---|
| Codex + Skills | C1 | Run 2 only | 2 | High | 100.0% | 0.0% |
| Codex + Skills | C5 | Run 2 only | 54 | High | 100.0% | 0.0% |
| DeerFlow 2 | C2 | Run 1 only | 5 | Low | 20.0% | 0.0% |
| DeerFlow 2 | C2 | Run 2 only | 3 | High | 100.0% | 0.0% |
| DeerFlow 2 | C3 | Run 1 only | 67 | Low | 0.0% | 0.0% |
| DeerFlow 2 | C3 | Run 2 only | 4 | High | 100.0% | 0.0% |
| DeerFlow 2 | C4 | Run 1 only | 234 | Low-Medium | 41.9% | 0.0% |
| DeerFlow 2 | C4 | Run 2 only | 221 | High | 100.0% | 0.0% |
| DeerFlow 2 | C5 | Run 1 only | 264 | Low | 1.5% | 0.0% |
| DeerFlow 2 | C5 | Run 2 only | 41 | High | 97.6% | 0.0% |
| DeerFlow 2 | C6 | Run 1 only | 3235 | Low | 0.2% | 58.9% |
| DeerFlow 2 | C6 | Run 2 only | 246 | High | 97.2% | 0.0% |
| Biomni | C1 | Run 1 only | 37 | Low | 10.8% | 2.7% |
| Biomni | C1 | Run 2 only | 2 | Low | 0.0% | 100.0% |
| Biomni | C2 | Run 1 only | 580 | Low | 20.9% | 8.4% |
| Biomni | C2 | Run 2 only | 1102 | Low | 7.7% | 90.8% |
| Biomni | C3 | Run 1 only | 1018 | Low | 0.0% | 2.2% |
| Biomni | C3 | Run 2 only | 11 | Low-Medium | 27.3% | 0.0% |
| Biomni | C4 | Run 1 only | 2083 | Low | 22.2% | 20.8% |
| Biomni | C4 | Run 2 only | 79 | Low | 2.5% | 6.3% |
| Biomni | C5 | Run 1 only | 1787 | Low | 15.1% | 0.0% |
| Biomni | C5 | Run 2 only | 543 | Low | 0.9% | 83.8% |
| Biomni | C6 | Run 1 only | 6182 | Low | 1.8% | 40.9% |
| Biomni | C6 | Run 2 only | 504 | Low | 48.8% | 47.8% |
